# Case report: intermittent fasting and probiotic yogurt consumption are associated with reduction of serum alpha-N-acetylgalactosaminidase and increased urinary excretion of lipophilic toxicants

**DOI:** 10.1101/177774

**Authors:** Jerry Blythe, Marco Ruggiero, Stefania Pacini

## Abstract

In this study, we describe the changes associated with three months of intermittent fasting and probiotic yogurt consumption in a 72-year-old marathon runner with chronic lymphocytic leukemia for a number of years. Serum alpha-N-acetylgalactosaminidase (nagalase), a marker of inflammation and cancer cell proliferation, was significantly decreased at the end of a three-month observation. These results are consistent with immune modulating properties of certain probiotics based on the fermentation of milk and colostrum. Urinary excretion of non-metal toxicants that accumulate in adipose tissue such as Perchlorate, N-acetyl(2-hydroxypropyl)cysteine (NAHP), 2,4-Dichlorophenoxyacetic acid, 3-Phenoxybenzoic acid (3PBA), N-acetyl phenyl cysteine (NAP), Phenylglycoxylic acid (PGO), Monoethylphthalate (MEP) and 2-Hydroxyisobutyric Acid (2HIB) was significantly increased. These results are consistent with the weight loss (5 Kg) associated with intermittent fasting and with the known features of probiotics as detoxification tools. Consistent with certain toxicants acting as endocrine disruptors, we observed an increased elimination of toxicants and a 33% decrease of serum Thyroid Stimulating Hormone (TSH), suggesting a trend toward normalization of thyroid function. These results support the hypothesis that a combination of intermittent fasting with the consumption of specific probiotic yogurts may lead to immune modulation, detoxification and other improvements.

**Abbreviations:**

(NAHP): N-acetyl(2-hydroxypropyl)cysteine
(3PBA): 3-Phenoxybenzoic acid
(NAP): Nacetyl phenyl cysteine
(PGO): Phenylglycoxylic acid
(MEP): Monoethylphthalate
(2HIB): 2-Hydroxyisobutyric Acid
(TSH): Thyroid Stimulating Hormone
(Dr. JB): Dr. Jerry Blythe
(CLL): chronic lymphocytic leukemia
(IRB): Institutional Review Board
(GcMAF): Gc protein-derived Macrophage Activating Factor

## Introduction

A recent review of the metabolic effects of intermittent fasting published in the July 2017 Annual Review of Nutrition suggests that nutritional regimens may “offer promising nonpharmacological approaches to improving health at the population level, with multiple public health benefits”.^(1)^ This statement is consistent with the conclusions of a study published in 2013 describing how “Intermittent fasting is reported to improve the lipid profile; to decrease inflammatory responses, reflected by changes in serum adipokine levels; and to change the expression of genes related to inflammatory response and other factors”.^(2)^ The interest in intermittent fasting as a nutritional approach to weight loss and a number of health issues, is demonstrated by the number of peer-reviewed papers on this topic. A PubMed query for “intermittent fasting” performed in July 2017, yielded, yielded 755 results with a quasi-exponential increase in the number of papers in the past 5 years.

Intermittent fasting appears particularly efficient in achieving sustained weight loss and enhancing body composition by reducing fat mass.^(3)^ And, a reduction of fat mass is often further associated with a release of lipophilic toxicants that may have accumulated in adipocytes^(3)^; such a release may pose a health danger if it is not accompanied by appropriate detoxification procedures. Among the different detoxification approaches, interventions involving modulation of the microbiota appear particularly promising as they target a system, in this instance, the microbiota that is involved in all aspects of human physiology. Such interventions have received a great deal of attention in the past few years.^(4,5)^

Thus, it has been known for more than 10 years that probiotic microbial strains have properties that enable them to bind metals and toxins from food and water; interestingly, binding of toxicants by probiotics appears to be instantaneous, thus leading to the proposal of exploiting these features for decontamination in food and intestinal models.^(6)^

Based on these premises, Dr. JB, a retired Medical Doctor, embarked on a three-month experience of intermittent fasting and probiotic yogurt consumption with the goal of weight loss and detoxification. In this study, we describe the changes associated with such a nutritional approach.

## Subject and methods

Dr. JB was born through Cesarean section in 1945, was fed with formulas, and suffered several early childhood infections, including pneumonia at the age of 5, treated with chloromycetin (one of the first commercial uses of chloramphenicol was chloromycetin in 1949). The drug is seldom used today because of potential toxicity and bone marrow depression. At the age of 6, he developed a passion for running that continues to this day. He kept logs of his mileage and calculates that he has run approximately 112,000 miles (180,246 Km) including a number of marathons; his remarkable achievements have recently been described in the journal, The Socionomist.^(7)^

He has been healthy and energetic throughout his adult life without illness other than Typhoid Fever acquired outside the United States in 1970 and treated with chloramphenicol. His bone marrow suffered a residual leukopenia and lymphocytosis which appear to have not affected his daily activities. In 2012, he was diagnosed with chronic lymphocytic leukemia (CLL) at an Indianapolis Cancer Center, and found to have a 13ql4.3 deletion. Retrospective study of blood analyses, however, suggests that the condition had been evolving over a number of years and diagnosable as CLL since 2008 when for the first time he developed a leukocytosis along with his lymphocytosis. After careful consideration, Dr. JB opted for complementary/alternative approaches for his condition with particular focus on healthy nutrition and exercise, avoiding refined sugars, alcohol, carbonated drinks (sodas), cookies, pies, candy or cakes; he maintained his consistent training program that he is still implementing, running on average 6 miles per day at a comfortable pace.

In March 2017, JB embarked on an intermittent fasting program patterned on the weight loss program described by He *et al.*^(3)^ Shortly thereafter, he introduced daily consumption of 100 ml of probiotic yogurt aimed at supporting the immune system and helping detoxification, choosing a product whose properties have been previously described.^(8,9)^

Blood and urine analyses were performed immediately before and after the three-month experience. Analysis of serum alpha-N-acetylgalactosaminidase (nagalase) was performed at the European Laboratory of Nutrients, (Bunnik, The Netherlands); the results are expressed as Units = nmol/min/mg. Blood analyses were performed at Lab Corp, Laboratory Corporation of America. Analyses of toxicants in early morning urine samples were performed at The Great Plains Laboratory Inc. (Lenexa, KS, USA) and are expressed as μg/g creatinine. Copies of the original records are conserved at the office of Dr. JB.

Since this is a single case report that does not produce generalizable knowledge, nor is it an investigation of an FDA regulated product, Institutional Review Board (IRB) review is not required for this activity.^(10)^ Likewise, since this case report described the experience of one of the Authors, Dr. JB, MD, the concept of “informed consent” is implicit in the authorship of the study.

## Results

After three months of intermittent fasting and probiotic yogurt consumption, Dr. JB’s body weight decreased from 70.7 to 66.2 Kg (6.4%). Serum nagalase values significantly decreased from 1.92 to 1.06 U (44.8%), a value that is very close to the upper normal value set by the laboratory (0.95). Serum TSH values also significantly decreased from 7.83 to 5.2 μlU/ml (33.6%). Leucocyte count decreased from 58 to 54.9 (cells × 10^3^/μL) (5.3%) and the percentage of lymphocytes decreased from 94 to 90% (4.4%). The number of platelets favorably increased from 93 to 125 (cells × 10^3^/μL) (34.4%).

Concomitant with these changes, urinary excretion of a number of lipophilic toxicants significantly increased (Table 1).

**Table 1.** Non-metal toxicants in urine.

| Compound | Before | After |
| --- | --- | --- |
| Monoethylphthalate (MEP) | 1,925 | 3,353 |
| 2-Hydroxyisobutyric acid (2HIB) | 4,364 | 7,349 |
| Phenylglyoxylic acid (PGO) | 36 | 184 |
| Perchlorate | 3.5 | 13 |
| N-acetyl phenyl cysteine (NAP) | N.D. | 0.11 |
| N-acetyl(2-hydroxypropyl)cysteine (NAHP) | 70 | 135 |
| 2,4-Dichlorophenoxyacetic acid | 0.18 | 1.7 |
| 3-Phenoxybenzoic acid (3PBA) | N.D. | 0.27 |

*Values are expressed as $\mu\text{g/g}$ creatinine; the first term (before) refers to the value observed before intermittent fasting and probiotic yogurt consumption; the second term (after) refers to the value observed after the three-month experience.*

It is worth noticing that the toxicants whose excretion was increased concomitantly with implementation of intermittent fasting and probiotic yogurt consumption, are among the most common environmental pollutants and potential sources of anthropogenic disturbance. It may be argued that elevated initial levels of phthalates were associated with Dr. JB’s consistent running and consumption of water from years of soft, plastic water bottles possibly containing phthalates. Interpretation of initial elevated level of 2HIB is more complex. Thus, 2HIB is formed endogenously as a product of branched-chain amino acid degradation in patients with ketoacidosis.^(11)^ However, 2HIB is also a metabolite of gasoline additives^(12)^, and elevated levels of 2HIB can indicate exposure to environmental pollution, exposure that could be associated with activities such as running in urban environments. Since Dr. JB had no clinical or laboratory signs of ketoacidosis, we interpret the elevated levels of 2HIB as consequence of environmental exposure.

## Discussion

The changes observed during a three-month experience of intermittent fasting and probiotic yogurt consumption seem to support the hypothesis that such a nutritional approach provides health benefits ranging from weight loss to detoxification and immune system support.

Decrease of nagalase can be interpreted as the result of modulatory activities of the probiotic yogurt on the immune system with particular reference to macrophage activation.^(8)^ In addition to macrophage stimulation, it is known that fermentation of milk leads to formation of immune-modulatory and anti-oxidant molecules that may play an additional role in decreasing the inflammatory status responsible for the elevated nagalase levels.^(13)^ Also, the probiotic yogurt used in this study contains bovine colostrum, a supplement known to have intrinsic immune-modulating properties^(14)^ and to be rich in vitamin D-binding protein that is the precursor of Gc protein-derived Macrophage Activating Factor (GcMAF).^(15)^ It has been proposed that nagalase is a marker of cancer cell proliferation ^(16)^ as well as of systemic inflammation.^(17)^ Therefore, the observed decrease of serum nagalase activity, together with the decrease of leucocytes and lymphocytes (leucocytes decreased from 58 to 54.9 cells × 10^3^/μL); lymphocytes from 94 to 90%) accompanied by an increase of platelets (from 93 to 125 cells × 10^3^/μL), may reflect a positive impact of the nutritional approach described herein on the underlying pathologic condition of Dr. JB. We attribute these effects to the ability of the probiotic yogurt used in this study to stimulate and balance macrophage function as we demonstrated since 2011.^(8,9)^

Such a positive impact may also be consistent with the observed decrease of serum TSH values. We interpret this change as a consequence of the increased excretion of toxicants known to act as endocrine disruptors with particular reference to thyroid function. For example, phthalates bind to thyroid hormone receptors altering the signaling of thyroid hormones ^(18)^, and perchlorate interferes with iodine uptake with resulting increased production of TSH as compensatory mechanism.^(19)^ Thus, it can be hypothesized that increased elimination of phthalates, perchlorate and other endocrine disruptors may be responsible for the trend toward normalization of thyroid function.

The observed increased urinary excretion of lipophilic toxicants is consistent with recent report describing how such a nutritional plan followed by Dr. JB is associated with an increase of serum concentrations of lipophilic pollutants such as polychlorinated biphenyls, possibly mobilized from adipose tissue.^(3)^ We attribute the increased urinary excretion of lipophilic chemicals to the effects of the cleansing procedures associated with the plan described in He *et al.*^(3)^ as well as to the known detoxifying effect of probiotics. For example, *Lactobacillus acidophilus*, a component of the probiotic yogurt used in this study, is endowed with antioxidant effects, decreasing malondialdehyde and increasing the levels of the antioxidants, glutathione reductase, superoxide dismutase, and glutathione peroxidase.^(20)^ Other probiotic strains provide significant protection against metal toxicity by decreasing the level of toxicants in the liver and kidney, and by preventing alterations in the levels of glutathione peroxidase and superoxide dismutase.^(21)^ Another mechanism of action responsible for the detoxifying effects attributable to probiotic yogurt consumption lays in the observation that lactic acid bacterial strains remove toxins from liquid media by physical binding^(22)^, and protect against toxins such as polycyclic aromatic hydrocarbons, heterocyclic aromatic amines, amino acid pyrolysates and mycotoxins.^(23)^ In addition, it has been recently demonstrated that probiotics such as those represented in the probiotic yogurt used in this study may improve kidney function thus contributing to the overall detoxifying effect.^(24)^

The authors wish to emphasize that the data in this case report are from one individual, and not part of a larger study. Like Aronson who encourages reports of single cases in medicine^(25)^, the authors believe the history and observations may benefit those addressing the impact of environment on health. The case of Dr. JB is rather peculiar for several reasons ranging from the intense training program to age and underlying chronic condition and, therefore, the results observed in this report cannot be generalized. Further studies are warranted to assess whether nutritional programs similar to the one here described will yield similar results in subjects of different age or with different conditions. For example, considering that environmental exposure to organic pollutants may play a significant role in autism^(26)^, it will be interesting to determine whether autistic children may benefit from the detoxification approach described in this report.

## Conclusions

The experience of Dr. JB suggests that intermittent fasting associated with probiotic yogurt consumption may lead to body weight reduction, increased excretion of toxicants and modulation of the immune system with overall positive effects on health.

## Acknowledgements

The Authors wish to thank Ms. Michelle Kriegel who introduced Dr. Blythe to the nutritional plan described in this study, and Mr. Peter Greenlaw, author of books on healthy nutrition, who connected him with Drs. Ruggiero and Pacini after completion of the three-month experience.

## Authors’ Contributions

Jerry Blythe, MD: performed the experience described in this study, provided critical input and assisted in revising and improving the paper.

Marco Ruggiero, MD, PhD and Stefania Pacini, MD, PhD: wrote the first draft of this paper, provided critical input and assisted in revising and improving the paper.

## Ethics

This article is original and contains material that has not been submitted or published in any other scientific journal. A draft manuscript has been archived in biorxiv, the pre-print server for biology (doi: https://doi.org/10.1101/177774).

